## Supplementary material for "Bactabolize: A tool for high-throughput generation of bacterial strain-specific metabolic models": Response to reviewer

Response to reviewers

### Reviewer #1

1. Overall, this is a useful tool, the data is well-presented, and the paper is well-written. However, the predictions are only compared with two existing reconstruction tools though more have been recently published
   - The aim of this work was to facilitate high-throughput generation of strain-specific metabolic models e.g. at the scale of 100s -1000s as indicated throughput the introduction (see lines 74-82, 91-94), and therefore we only compared tools which were capable of high-throughput analysis via command line and excluded others (e.g. merlin). We have now tested this against the other recent command line tool, gapseq which had escaped our gaze. Thank you for bringing this to our attention. Additionally, we have included KBase (ModelSEED, a web-based app that does not support high-throughput analysis) to allow for readers to interpret the results in the context of community standard approaches, since KBase is a popular tool.
   - We have added an explicit statement about the choice of approaches, now at lines 194-199 as follows:
     “We compared the output and performance of Bactabolize to the two previously published tools that can support high-throughput analyses i.e. CarveMe (30) and gapseq (31). To aid interpretation in the context of community standard approaches, we also include a comparison to the popular web-based reconstruction tool, KBase (ModelSEED), and a manually curated metabolic reconstruction of K. pneumoniae strain KPPR1 (also known as VK055 and ATCC 43816, metabolic model named iKp1289) (15).”
   - The methods section was updated accordingly (now lines 552-558):

“A draft model was generated using gapseq version 1.2 with the ‘doall’ command using the unannotated genome (as gapseq does not take annotated input files). Gap-filling was subsequently performed using the ‘fill’ command and a custom M9 media file to match the nutrient list found in Bactabolize (https://github.com/kelwyres/Bactabolize/blob/main/data/media_definitions/m9_media.json).

Finally, a draft model was constructed using the annotated genbank K. pneumoniae KPPR1 file and the KBase narrative (15) (https://narrative.kbase.us/narrative/ws.14145.obj.1).”

- The results section (now lines 193-308) and Figure 2 have also been updated / restructured to reflect the new analyses, and include a comparison of the relative compute times for the construction of models (lines 281-291) as follows:

“While model features and accuracy are essential metrics for comparison, computation time is also a key consideration for high-throughput analyses. We recorded the time required for each tool to build draft models for 10 of the completed KpSC genomes used in the quality control framework (see below) on a high-performance computing cluster (Intel Xeon Gold 6150 CPU @ 2.70GHz and 155 GB of requested memory on a CentOS Linux release 7.9.2009 environment. CarveMe KpSC pan was the fastest with a mean of 20.04 (range 19.90 - 20.18) seconds, followed by CarveMe universal at 30.28 (range 29.20 - 31.80) seconds, then Bactabolize KpSC pan at 98.05 (range 92.19 - 100.4) seconds. KBase took 183.50 (range 120.00 - 338.00) seconds per genome via batch analysis, including genome upload time and queuing. gapseq took 5.46 (range 4.55 - 6.28) hours to produce draft models (not including the required gap-filling), consistent with previous reports (37).”

- Finally, the whole discussion has been updated and substantially restructured (lines 472-474, 475-493, 494-512). Specific mentions to the new analyses are at lines:

472-474: “Consistent with this assertion, our draft KPPR1 model constructed with KBase (without manual curation) was an outlier in terms of the very low number of genes, reactions and metabolites that were included.”

2. My understanding is that the tool requires a set of reference reconstructions for other strains of the target species. If no reference reconstruction is available for another strain of the target species, can this species not be reconstructed?
   - Any input reference can be used to generate models however, single strain models matching the target species, or ideally a species-specific pan-reference, are recommended for best results. We have added a discussion on these points at lines 128-133:

3. How do the reconstructions generated by Bactabolize compare to those generated by other reconstruction tools besides CarveMe and ModelSEED, e.g., gapseq (Zimmermann et al, Genome Biology 2021. 22:81) or merlin (Capela et al, Nucleic Acids Res 2022, 50(11):6052-6066?
   - See response to rev 1 point 1.
4. How are the accuracy, specificity, and sensitivity of the pan-models calculated? Is the compared experimental data on the species level?
   - We used the pan-model as a reference from which we generated a strain-specific model for *K. pneumoniae* KPPR1 (using Bactabolize and CarveMe). This strain-specific metabolic model was then used to simulate growth phenotypes and compared to published experimental data for KPPR1. This was described in the methods section, including the calculations for the metrics (lines 589-593); however, we have also expanded the description within the results section to clarify the approach (lines 201-209):

“De novo draft models for strain KPPR1 were built using; i) Bactabolize with the KpSC pan v1 reference; ii) CarveMe, with its universal reference model (CarveMe universal); iii) CarveMe, with KpSC-pan v1 reference (CarveMe KpSC pan); iv) gapseq; and v) KBase (ModelSEED). ….. Subsequently, each model was used to predict growth phenotypes; i) in M9 minimal media with different sole sources of carbon, nitrogen, phosphorus and sulfur; and ii) for all possible single gene knockouts in LB under aerobic conditions. The predicted phenotypes were compared directly to the published phenotype data.” [Note the published data are cited in the previous manuscript sentence, not shown here].

5. The link https://github.com/rrwick/GFA-dead-end-counter, in line 286 does not work.
   - Link regenerated – now at line 451-452 and 604

### Reviewer #2

1. KpSC pan-metabolic reference model is provided. Are they required as input for Bactabolize? Are the gene, metabolite information open accessible by users?
   - See response to reviewer 1 point 2 above and;
   - All data for the KpSC pan-model described in this work are accessible in the model files and amino acid + nucleotide files + data table at <https://github.com/kelwyres/KpSC-pan-metabolic-model>. This is also linked in the manuscript at line 631 and in the Data availability statement at line 661.
2. In the results section "description of Bactabolize", the authors present technical details on how to generate a metabolic model. For the input and output, please provide concrete examples to show the functionality of Bactabolize.
   - Detailed instructions, example code and example input/output files are available via the Bactabolize GitHub repository: <https://github.com/kelwyres/Bactabolize>.

Instructions and example code can be found on the wiki: <https://github.com/kelwyres/Bactabolize/wiki>

Test data and example files are at: <https://github.com/kelwyres/Bactabolize/tree/main/data/test_data>

The Github repository is linked in the manuscript at lines 95, 124, 552, and 667, and we have added a further reference at line 124, which mentions the example code/data:

“Full documentation, including example code and test data are available at the Bactabolize code repository (https://github.com/kelwyres/Bactabolize).”

3. To generate metabolic models, the authors present comparison results with other methods. However, the authors only present the numbers in genes, metabolites and substrates. Since the interactions between gene, metabolite, and substrate are also critical, if possible, please provide the coverage details about these interactions. Venn diagram is recommended to compare these coverage differences.
   - Two additional supplementary figures have been generated (Figures S5 and 6) showing Venn diagrams of metabolites and reactions for the high-throughput analysis approaches that are most relevant to this work (see also response to rev 1, point 1). These are discussed at lines 224-237:

4. Are quality control and gap-filling needed to be processed when constructing a new metabolic model?

Our goal here was to implement an approach to support high-throughput analyses (see response to rev 1 point 1), including leveraging draft genome assemblies as the bases for the construction of strain-specific metabolic models. As part of this work, we have described a robust quality control (QC) framework for screening draft *K. pneumoniae* genomes i.e. to identify genome assemblies that should not be used. We developed this framework by comparison to models generated for matched completed genomes. Our analyses demonstrate the importance of applying QC to the input draft genome assemblies. When appropriate QC is applied to the input genomes, the resultant draft models show a high degree of completeness compared to the matched models derived from complete genomes. The draft models can also be used to simulate growth phenotypes with high accuracy as compared to those simulated for the matched complete genome models.

No specific QC was applied to the draft models themselves, other than confirmation of positive growth prediction in m9 minimal media plus glucose (which is expected to support growth of all K. pneumoniae). In cases where the input assembly passed our QC criteria but the resultant model was unable to simulate growth in m9 minimal media plus glucose, gap-filling may be optionally applied. Again, by comparison to the simulated phenotypes from matched complete genome models, we show that these gap-filled draft models can produce accurate phenotype predictions. See lines 396-404:
“Of the 901 draft genome assemblies which passed our QC criteria (≤200 assembly graph dead ends), 23 of the resulting draft models failed to simulate growth in M9 minimal media with glucose (despite capturing ≥99% of the genes and reactions in the corresponding complete models). It is expected that all KpSC models should be able to simulate growth on M9 media with glucose as a sole carbon source, as this central metabolism is universal amongst KpSC. To replace missing, critical reactions required for growth on M9 with glucose, we investigated model gap-filling using the patch_model command of Bactabolize. We then assessed the accuracy of the gap-filled models for prediction of growth on the full range of substrates, as compared to the predictions from the corresponding complete models.”

Lines 409-413: “Substrate usage predictions from the 21 successfully gap-filled models were highly accurate, with 18/21 having a prediction concordance of ≥99% across all 846 growth conditions (12/21 had 100% concordance) (Figure S9). We therefore conclude that models generated for genome assemblies passing our QC criteria, which have been gap-filled to successfully simulate growth on minimal media plus glucose, are suitable for the prediction of growth across a range of substrates.”

5. Are there any visualization results to check the status of the generated draft model?
   - No. This is a tool for large-scale and rapid production of metabolic models, and phenotype prediction and we have not included visualisation tools. Third party tools are available e.g. <https://fluxer.umbc.edu/>. We do provide optional generation of MEMOTE reports at lines 136-138:
     “Draft genome-scale metabolic models are output in both SMBL v3.1 (41) and JSON formats (one pair of files for each independent strain-specific model), along with an optional MEMOTE quality report (42)”.

### Reviewer #3

1. The justification and evaluation of the generated models are inadequate and one-dimensional. The authors only focus on statistics such as the number of reactions and genes in the models, which does not accurately depict the completeness of the model.
   - The reviewer has misunderstood how we have used ‘completeness’ in this manuscript. In the section describing our novel QC framework, we use this term to refer to the relative completeness of draft models generated from draft genome assemblies as compared to curated models generated from complete genome assemblies for the same strains. The latter were considered as the ‘complete’ models for this purpose. We are not referring to any measure of network or metabolic pathway completeness. We specifically refer to gene and reaction capture compared to the ‘complete’ models because these features directly reflect the problem we are trying to address i.e. that draft genome assemblies may not contain the complete set of genes that are truly present in the underlying genome. We have updated the manuscript text to further clarify the problem we aim to address in this section and justify the use of gene and reaction capture metrics:

Lines 310-319: “There are now thousands of bacterial genomes available in public databases, the majority of which are in draft form, comprising 10s to 1000s of assembly contigs. This fragmentation of the genome is caused by repetitive sequences that cannot be resolved by the assembly algorithm and/or sequence drop-out. The latter can result in the loss of genetic information such that some portion of genes present in the underlying genome are lost from the genome assembly (either completely or partially). This in turn, poses a limitation for the reconstruction of metabolic models using these assemblies, since most published approaches use sequence searches to predict the presence/absence of genes and their associated enzymatic reactions. Therefore, if we are to use public genome data for high-throughput metabolic modelling studies, it is essential to evaluate the expected model accuracies and understand the minimum input genome quality requirements.”

The biological accuracy of the curated ‘complete’ models has been described previously, and this is now noted in the text at lines 320-324:

“Here we performed a systematic analysis leveraging our published curated KpSC models (n=37, (14)), which were generated using completed genome sequences and were therefore considered to represent ‘complete’ models for which the underlying genome sequence contains all genes that are truly present in the genome (note the biological accuracy of these models was reported previously (14) and is not the subject of the current study).”

- Throughput the manuscript we not only compare models in terms of the numbers of genes and reactions, but through comparison of binary growth predictions. Specifically, in the **Performance Comparison** section (Bactabolize vs other approaches) we use comparison of predicted to experimental phenotypes for strain KPPR1 (see response to rev 1 point 4 for details). In the **QC Framework** section we compare the predictions derived from draft models generated from draft genome assemblies to those derived from the matched ‘complete’ models, and report the concordance as a measure of impact of input assembly quality (lines 309-394). In the final results section (**Predictive accuracy of draft models**), we generate 10 additional models and compare the growth predictions to matched experimental data (lines 414-433). We view these phenotype prediction comparisons as the ultimate measure of ‘completeness’ with which to assess our models, because these data have direct biological meaning.

2. The authors have not provided evidence or discussion on the accuracy of any metabolic fluxes, which are considered to be crucial for reconstructing metabolic models. Additionally, the authors have not mentioned the importance of non-growth associated maintenance and the criticality of biomass composition analysis, both of which significantly determine the fluxes in the system.
   - We acknowledge the importance of flux calculations and accurate biomass compositions when using genome-scale models to quantitatively predict growth rates. However, at this stage, the reconstructions developed using Bactabolize are intended for *binary* predictions and comparisons of growth capabilities on various substrates. The accuracies we report are based on measures of network completion (presence/absence of relevant reactions leading to growth or no-growth phenotypes) rather than specific growth rates. Thus, the models generated by Bactabolize can be used to explore diversity at the strain level in terms of growth capabilities and can serve as a scaffold for building detailed (customized biomass), strain-specific models. Measuring biomass composition and metabolic flux analysis require significant experimental comparisons that are outside the scope of the current study but could be performed for target strains based on reconstructions developed using Bactabolize.
3. It would be interesting to compare the accuracy of the models generated using Bactabolize with those manually curated.
   - We did exactly this. We compared the manually curated model iKp1289 as part of our benchmarking. Lines 194 – 199:
     “*We compared the output and performance of Bactabolize to the two previously published tools that can support high-throughput analyses i.e. CarveMe (30) and gapseq (31). To aid interpretation in the context of community standard approaches, we also include a comparison to the popular web-based reconstruction tool, KBase (ModelSEED), and a manually curated metabolic reconstruction of K. pneumoniae strain KPPR1 (also known as VK055 and ATCC 43816, metabolic model named iKp1289) (15).*”

     Unfortunately, as far as we aware there are currently no other published manually curated models for strains with matched phenotype data that are also not included as part of our pan-reference model (the latter is a key point to ensure a fair comparison of models generated using our pan-reference vs those generated with a universal reference).
4. The authors have not provided evidence or discussion on the accuracy of any metabolic fluxes, which are considered to be crucial for reconstructing metabolic models.
   - See response to rev 3, point 2.
5. The justification regarding the completeness of the models requires further discussion.
   - See response to rev 3, point 1.
6. A detailed discussion on the importance of manually curated models would significantly enhance the quality of the manuscript.
   - This has been added at lines 458-474:

“Traditionally, genome-scale metabolic reconstruction approaches have relied upon significant manual curation efforts. While there will always remain a need for high quality curated models, such resource intensive approaches preclude their application at scale, and have therefore limited analyses to small numbers of individual strains (15, 16). However, automated reconstruction approaches can support the generation and comparison of multiple strain-specific draft models from which meaningful biological insights can be derived (61). Additionally, the quality of curated models is likely to vary depending on their age, level and type of curation, as well as the approach used for preliminary drafting. Indeed it is possible for automated approaches to outperform manually curated models; a draft model for K. pneumoniae KPPR1 generated using Bactabolize with the KpSC pan-v1 reference model outperformed the manually curated iKp1289 model representing the same strain (15). iKp1289 was published in 2017 (6 years prior to this study) and was initially drafted via the KBase pipeline (33), which uses RAST to annotate the sequences with Enzyme Commission numbers. It has been demonstrated several times that the Enzyme Commission scheme has systematic errors (62, 63), leading to a loss in accuracy when compared to the ortholog identification methods used by automated approaches. Consistent with this assertion, our draft KPPR1 model constructed with KBase (without manual curation) was an outlier in terms of the very low number of genes, reactions and metabolites that were included.”
